# Is physiological stress state reflected in acoustic structure of vocalizations? An experimental test in wild North American red squirrels

**DOI:** 10.1101/456830

**Authors:** Matt Sehrsweeney, David R. Wilson, Maggie Bain, Stan Boutin, Jeffrey E. Lane, Andrew G. McAdam, Ben Dantzer

**Affiliations:** Department of Psychology, University of Michigan, Ann Arbor, MI, USA; Department of Psychology, Memorial University, NL, Canada; Department of Integrative Biology, University of Guelph, Guelph, ON, Canada; Department of Biological Sciences, University of Alberta, Edmonton, AB, Canada; Department of Biology, University of Saskatchewan, Saskatoon, SK, Canada; Department of Ecology & Evolutionary Biology, University of Michigan, Ann Arbor, MI, USA

## Abstract

Acoustic signaling is an important means by which animals communicate both stable and labile characteristics. Although it is widely appreciated that vocalizations can convey information on labile state, such as fear and aggression, very few studies have experimentally examined the acoustic expression of short-term stress state. The transmission of such information about physiological state could have broad implications, potentially allowing other individuals to modify their behavior or life history traits in response to this public information. North American red squirrels (*Tamiasciurus hudsonicus*) produce vocalizations known as rattles that advertise territorial ownership. We examined the influence of changes in physiological stress state on rattle acoustic structure through the application of a stressor (trapping and handling the squirrels) and by provisioning squirrels with exogenous glucocorticoids (GCs). We characterized the acoustic structure of rattles emitted by these squirrels by measuring rattle duration, mean frequency, and entropy. Our results provide mixed evidence that rattles show a “stress signature”. When squirrels were trapped and handled, they produced rattles that were longer in duration with a higher frequency and increased entropy. However, squirrels that were administered exogenous GCs had similar rattle duration, frequency, and entropy as squirrels that received control treatments and unmanipulated (unfed) squirrels. Our results indicate that short-term stress does affect the acoustic structure of vocalizations, but elevated circulating GC levels are not solely responsible for such changes.

## Introduction

Acoustic communication is a critical means by which information is transferred within and among animal species. Vocalizations can convey stable information on various characteristics of signalers, such as individual identity (Beer, 1970; Beecher, 1989; Blumstein and Munos, 2005), body weight and size (Fitch 1997; Bee et al. 1999; Reby and McComb, 2001; Blumstein and Munos, 2005; Koren and Geffen, 2009), sex (Ey et al., 2007; Blumstein and Munos, 2005), and social rank (Clark, 1993; Koren et al., 2008; Muller et al., 2004; Terleph et al., 2016; Yosida and Okanoya, 2009), and they are often encoded with several layers of information (Koren and Geffen 2009). Communicating this information is consequential for both signalers and receivers, serving a wide array of functions, from attracting mates (Andersson 1994) to reducing conflict and maintaining affiliations in social groups (Masataka and Symes, 1986; Digweed, et al., 2007; Soltis et al., 2005a).

Vocalizations can also contain information on labile, or short-term, traits, such as short-term stress or perhaps the changes in glucocorticoids (GCs) that are released in response to a short-term stressor. Stress is associated with elevated glucocorticoid levels and is known to influence the acoustic structure of vocalizations in a number of species (Manser, 2001; Wilson and Evans, 2012; Sacchi et al., 2002; Slocombe et al., 2009). Motivation-structural rules make predictions about the characteristics of vocalizations produced in high-stress contexts: hostile vocalizations tend to be lower in frequency and noisier (highly entropic), and fearful vocalizations tend to be higher in frequency and more tonal (Morton 1977). Although some studies have found empirical support for these rules, others have found inconsistencies. For example, vocalizations associated with fear often fail to consistently conform to these motivation-structural rules, and are often highly entropic (Morton 1977; August and Anderson 1987). The effects of short-term stress on vocalization structure are thus difficult to generalize.

Possible mechanisms by which short-term stress affects the acoustic structure of vocalizations have also been investigated. Short-term stress in known to induce a withdrawal of vagal output from motor neurons in the nucleus ambiguus that causes vocal muscles to tighten, resulting in an increase in vocalization pitch (Porges, 1995). In addition, entropy of vocalizations tends to increase with short-term stress because the oscillation of the laryngeal folds, which govern volume, reaches its maximum amplitude but subglottal pressure continues to increase, resulting in vocal non-linearities, including noise (Fitch et al., 2002).

Although many studies have examined the structure of vocalizations produced in high stress situations, they have concentrated primarily on vocalizations produced in just a few contexts, and most of them have been observational. Most studies have focused on social contexts, including calls produced by victims in agonistic encounters between social group members (Morton, 1977), alarm calls (Zuberbuhler, 2009), separation between mothers and their young, and between social group members (Biben et al., 1986; Ehret, 2005; Bayart et al. 1990; Rendall, 2003). Other research has centered on begging calls (Sacchi et al., 2002; Perez et al., 2016) and distress screams produced by individuals in imminent danger of predation or of being seized by a predator, which likely function to solicit intervention from another animal capable of interfering (Hogstedt, 1982; Lingle et al., 2007; Blumstein et al., 2008).

Very few studies have experimentally examined the influence of stress or changes in glucocorticoids on vocalization structure. One notable exception is Perez et al. (2012), who assessed the effects of GCs on the acoustic structure of Zebra Finch (*Taeniopygia guttata*) vocalizations. Their experiment included two stress treatments: social isolation, and treatment with exogenous GCs, and they found that both types of stress significantly altered vocalization features. Compared to untreated individuals, finches in both treatment groups emitted vocalizations of higher frequency than finches in the control group (Perez et al., 2012).

The literature on the influence of stress on vocalizations skews heavily towards group-living species and focuses primarily on just a few contexts in which stress occurs; far less is known about the relationship between stress and vocalization structure in solitary species, despite the fact that many regularly produce vocalizations in short term stress inducing situations (Hogstedt, 1982). Furthermore, few studies have experimentally examined this relationship, leaving a gap in our understanding of the mechanism by which stress may influence acoustic structure. We examined how a short-term stress (resulting from trapping and handling) and administration of exogenous GCs affected the territorial vocalizations of solitary, territorial North American red squirrels (*Tamiasciurus hudsonicus)*. Red squirrels defend discrete territories throughout the year, and produce vocalizations called “rattles” that advertise territorial ownership (Smith, 1968), which deters intruders (Siracusa et al., 2017). At the center of each territory is a “midden,” a network of underground tunnels that serves as storage space for white spruce (*Picea glauca*) cones that compose 50-80% of a squirrel’s annual diet (Donald et al. 2011; Fletcher et al., 2013). Overwinter survivorship without a midden is near zero (Larsen and Boutin, 1994). Successful defense of a territory against pilferage from the midden, therefore, represents an important component of overwinter survival for a red squirrel.

Red squirrel rattles contain stable information on individual identity (Digweed et al. 2012; Wilson et al., 2015), and receivers discern encoded kinship information, though this may be context-dependent (Wilson et al., 2015; Shonfield et al., 2017). In a playback experiment, focal squirrels only differentiated between the rattles of kin and non-kin when the playback rattles used were emitted by squirrels that had just been live-trapped and handled (henceforth, “post-trap rattles”), suggesting that post-trap rattles, produced in a high-stress state, are structurally distinct from rattles collected opportunistically (Shonfield et al., 2017). Thus, preliminary evidence suggests that rattles may encode information about stress state.

To test this directly, we conducted a two-part study to examine the relationship between stress state and rattle acoustic structure. In the first experiment, we recorded rattles of wild red squirrels after they were live-trapped and handled and compared these to rattles recorded opportunistically, without provocation, from squirrels moving freely around their territories. Previous studies verified this method of inducing stress: squirrels exhibit a substantial increase in circulating GC levels minutes after entering a trap and during handling (Bosson et al., 2012; van Kesteren et al., submitted).

To identify if elevated circulating GCs are part of the mechanism by which a short term stressor (such as capture and handling) alters rattle acoustic structure, we conducted a second experiment where we treated squirrels with GCs (dissolved in a small amount of food) and compared their rattles to those of squirrels in a control group (provided with the same amount of food but without GCs) and an unmanipulated group (provided with no food or GCs). A previous study showed that in GC-treated squirrels, plasma GCs rose quickly after treatment and then slowly declined over the ensuing 12 hours (van Kesteren et al., submitted).

In the first experiment, we predicted that if rattles do encode information about stress state, post-trap rattles would be structurally distinct from opportunistic rattles. Based on the results of Perez et al.’s (2012) Zebra Finch experiments, we predicted that post-trap rattles would be higher in frequency. In the second experiment, we predicted that if GCs are the mechanism by which short term stress alters rattle acoustic structure, rattles emitted shortly after treatment with exogenous GCs would exhibit the same structural distinctions as post-trap rattles when compared with rattles produced prior to treatment and rattles produced by control and unmanipulated squirrels over the same period of time. We expected these structural distinctions to be graded, peaking shortly after treatment and then declining as a function of time since consumption of treatment mirroring the peak and decline of circulating GC levels following treatment.

## Methods

### Study Site and Species

This study was part of the Kluane Red Squirrel Project, a long-term study of a wild population of red squirrels that has been tracked continuously since 1987 (McAdam et al., 2007), within Champagne and Aishihik First Nations traditional territory in the southwestern Yukon (61° N, 138° W). The habitat is an open boreal forest dominated by white spruce trees (*Picea glacua*; Krebs et al. 2001). All squirrels were marked individually with ear tags with distinct alphanumeric combinations (Monel #1; National Band and Tag, Newport, KY, USA), and wires in unique color combinations were threaded through the ear tags to allow for individual identification from a distance. We live trapped squirrels periodically to track female reproductive state and territorial ownership using tomahawk traps (Tomahawk Live Trap Company, Tomahawk, WI, USA) baited with peanut butter (McAdam et al. 2007).

### Experiment 1 Field Methods

We collected rattles from squirrels across four study areas between April and August in six separate years from 2005 and 2017 (Table 1). In the capture-induced stress experiment, we compared the structure of rattles collected opportunistically to rattles collected shortly after a squirrel was trapped, handled, and released. We collected rattles for this experiment using a Marantz digital recorder (model PMD 660; 44.1 kHz sampling rate; 16-bit amplitude encoding; WAVE format) and a shotgun audio recorder (Sennheiser, model ME66 with K6 power supply; 40-20,000 Hz frequency response (± 2.5 dB); super-cardioid polar pattern). To collect opportunistic rattles, we stood on a squirrel’s midden at a distance of no greater than 5 m from the squirrel until it produced a rattle. Red squirrels rattle spontaneously and in response to detection of conspecifics (Smith 1978), but we cannot rule out the possibility that the rattles were elicited by the person recording. A post-trap rattle was the first rattle emitted upon the squirrel’s release from a handling bag after spending time in a trap (within a minute of release). We did not record the exact amount of time a squirrel spent inside of a trap, but squirrels were in traps for no more than 120 min before they were released and a rattle was collected. As would be expected, squirrels exhibit a substantial increase in circulating GC levels minutes after entering a trap and during handling (Bosson et al., 2012; van Kesteren et al. submitted). In total, 351 rattles from 235 unique individuals (308 opportunistic rattles from 205 squirrels, 39 post-trap rattles from 30 squirrels) were collected and analyzed in the years 2005, 2006, 2009, 2015, and 2016. Of the 235 squirrels, 127 were male and 108 were female (Table 1). These rattles were part of a long-term dataset of rattles compiled by prior researchers with the Kluane Red Squirrel Project.

**Table 1:** Number of rattles collected by year, study grid, collection method, and date range. In parentheses, rattles are split up by sex - (male, female). For some squirrels, more than one rattle was collected.

| Year | Grid: AG | Grid: KL | Grid: SU | Grid: JO | Collection Method | Date Range |
| --- | --- | --- | --- | --- | --- | --- |
| 2005 | 0 | 2<br>(2,0) | 3<br>(1,2) | 0 | Opportunistic: 1<br>Post-trap: 4 | Jun 7 - Jul 31 2005 |
| 2006 | 0 | 113<br>(66,47) | 93<br>(43,50) | 0 | Opportunistic: 204<br>Post-trap: 2 | Jun 13 - Jul 14 2006 |
| 2009 | 30<br>(15,15) | 53<br>(26,27) | 8<br>(6,2) | 0 | Opportunistic: 54<br>Post-trap: 37 | Mar 26 - Jul 26 2009 |
| 2016 | 24<br>(12,12) | 25<br>(14,11) | 0 | 0 | Opportunistic: 49<br>Post-trap: 0 | Jun 6 - Aug 2 2016 |
| 2017 | 0 | 93<br>(93,0) | 22<br>(22,0) | 599<br>(599,0) | Zoom mic: 714<br>- Unmanipulated: 115<br>- Control: 367<br>- GC: 232 | Jun 2 - Aug 14 2017 |

### Experiment 2 Field Methods

In the second experiment, we assessed the influence of experimental increases in circulating GCs on rattle acoustic structure. We sought to track graded changes in rattle acoustic structure over an extended period of time induced by the GC treatment instead of a simpler pre/post treatment analysis. We compared the rattles of squirrels in three treatment groups, using an established protocol for oral administration of GCs. In the experimental group (n = 16), individuals were fed 8 g of peanut butter mixed with 2 g of wheat germ and 8 mg of cortisol (hydrocortisone, Sigma H004). This treatment causes a significant increase in circulating GCs (Dantzer et al 2013; van Kesteren et al submitted). Individuals in the control group (n = 16) were fed the same amount of peanut butter and wheat germ, with no cortisol added. Each squirrel in these two treatment groups was treated for one day (see details below). Lastly, we had an unmanipulated group of squirrels that were not fed or manipulated in any way (n = 23). Our treatment groups and our unmanipulated group of squirrels (living on a nearby study area) were comprised exclusively of male squirrels. However, no sex differences are known to exist in rattle acoustic structure (Wilson et al., 2015).

The morning of treatment, between 0730 and 1000 h, for each squirrel in the GC treated and control groups, we placed one treatment in a bucket hanging in a tree near the center of its midden. Pilferage from buckets was extremely low (van Kesteren et al, submitted), ensuring that treatments were eaten by the target squirrel, and not neighboring conspecifics or heterospecifics. We recorded the time each treatment was placed in each bucket and checked the buckets throughout the morning at a minimum of once every hour and maximum of every 45 min in order to determine the one-hour time frame in which the squirrel consumed its treatment. All treatments were consumed between 0830 and 1130. Eight squirrels (Control n = 4; GC, n = 4) did not consume their treatments by 11:30; these treatments were removed from the bucket and the squirrels were excluded from analyses. Two individuals (GCs n = 2) consumed their treatment over a period of several hours instead of consuming it within a one-hour time block. Because we sought to simulate short-term stress induced by a rapid elevation of circulating GC levels, these squirrels were excluded from analysis as well. Our final sample size was GC (n = 10), control (n = 12), and unmanipulated (n = 23).

We recorded rattles using stationary Zoom H2N Audio Recorders (Zoom Corporation, Tokyo, Japan) that were covered with windscreens and attached to 1.5 m stakes in the center of each squirrel’s midden. Because they are not weather-proof, we placed an umbrella 30 cm above each audio recorder to protect it from harsh weather conditions. We set the audio recorders in 44.1kHz/16bit WAVE format and recorded in 2-channel surround mode. We deployed the audio recorders between 1700 and 2200 h on the day before treatment so that they would collect “pre-treatment” rattles the following morning, prior to treatment. They recorded continuously until nightfall on the day of treatment, recording rattles of the target squirrel, neighboring individuals, and other ambient noise. Rattles recorded in the evening prior to treatment were excluded from analysis; thus, all rattles analyzed in this experiment were recorded on the day of treatment, between approximately 0600 and 2330 h. We chose this recording period because this recording window should have captured rattles at natural GC levels (pre-treatment rattles), during the post-treatment spike in circulating GC levels, and the ensuing decline. This is based upon our previous study showing that when squirrels are fed exogenous GCs, plasma cortisol concentrations spike shortly after treatment and decline over the ensuing 12 h (van Kesteren et al., submitted).

In order to analyze rattles recorded on stationary zoom recorders, we used Kaleidoscope software (version 4.3.2; Wildlife Acoustics, Inc., Maynard, MA, USA) to detect rattles in the recordings. Detection settings were: frequency range: 2000-13000 Hz; signal duration: 0.4-15 s; maximum intersyllable silence: 0.5 s; fast Fourier transform size: 512 points (corresponding to a temporal resolution of 6.33 ms and a frequency resolution of 86 Hz); distance setting: 2 (this value ensures that all detections are retained). Previous research using our same population, recording apparatus, and rattle extraction technique, and ground-truthed by comparing the results to those obtained by a human observing the squirrels being recorded, showed that our method detects 100% of a focal squirrel’s rattles (see Siracusa et al., submitted), but also detects non-rattles and the rattles of neighbors.

We used a previously developed a technique for distinguishing focal squirrel rattles from non-rattles and neighbor rattles (Siracusa et al., submitted). We first automatically analyzed the acoustic structure of every detection using the R package ‘Seewave’ (version 2.0.5; Sueur et al. 2008). Structural features included duration, root-mean-square amplitude, pulse rate, duty cycle, peak frequency, first energy quartile, skewness, centroid, and spectral flatness (see detailed definitions in Sueur et al. 2008 and Siracusa et al., submitted). Second, we used SPSS (software, version 24, IBM Corporation, Armonk, New York, USA) to apply a previously established linear discriminant function analysis model to the structural measurements of each detection. The model, which was developed during the same ground-truthing experiment described above, labeled each detection as ‘focal rattle,’ ‘neighbor rattle,’ or ‘non-rattle,’ and assigned a probability that the detection was a focal rattle. Third, we used Kaleidoscope to review spectrograms of all detections labeled ‘focal rattle’ that have an estimated probability of being a focal rattle of at least 0.99999. During this step, we removed any non-rattles that were included erroneously as focal rattles.

Our final dataset included 714 rattles from 45 focal squirrels (GC-fed = 232 rattles from 10 squirrels, control = 367 rattles from 12 squirrels, and unmanipulated = 115 rattles from 23 squirrels). Based on a cross-validated assessment of the accuracy of our approach (see details in Siracusa et al, submitted), 52% of all focal rattles should have been identified correctly as focal rattles (*i.e.,* 48% incorrectly classified as coming from a neighbour, and, therefore, excluded; false negative error rate = 48%), and 6% of the rattles labeled as focal rattles (after manually removing the non-rattles) should actually have been neighbor rattles (*i.e.,* false error rate of 6%). Therefore, although our final dataset included only half of all rattles produced by our focal squirrels during their 24-h trials, the vast majority of rattles that were included in the dataset were from the focal individual.

### Acoustic Analysis

We used Avisoft SASLab Pro software version 5.0 (Avisoft, 2015) to analyze the acoustic structure of rattles recorded in both experiments. The rattles were loaded into Avisoft, and for each rattle we generated a spectrogram (FFT size: 512, Window: Hamming, Temporal Resolution: 1.45 ms, Frequency Resolution: 86 Hz, Overlap: 87.5%) and the program extracted the acoustic parameters of interest (described below) using an existing protocol for rattle acoustic analysis, under our manual oversight. To characterize rattles, we measured three acoustic parameters: rattle duration, mean frequency (the frequency below which lies 50% of the energy of the signal, as measured from an averaged power spectrum of the entire signal), and entropy, a measure of noisiness of a signal. Because rattles are broadband and noisy signals, meaning that the majority of the energy in a call is dispersed across the frequency domain, mean frequency (the frequency below which lies 50% of the energy of the signal, measured from an averaged power spectrum of the entire signal) is a more appropriate measure of the frequency of the call than peak frequency. AviSoft measures Weiner Entropy (spectral flatness), calculated by dividing the geometric mean of the power spectrum by the arithmetic mean of the power spectrum, which ranges from 0 (pure tone) to 1 (white noise). These measurements were made using the ‘automatic parameter measurements’ feature of SASLab Pro to eliminate human bias in the measurements (settings: threshold −13 dB, hold time of 150 ms).

Because high frequencies attenuate more readily than low frequencies, entropy and mean frequency could, in theory, covary with recording distance. In the capture-induced stress experiment, a constant recording distance of approximately 5 meters was maintained for all recordings. In the GC induced stress experiment, in which rattles were recorded on stationary zoom microphones, to ensure that recording distance did not vary with time or treatment, we measured the signal-to-noise ratio of a subset of 140 rattles and found no significant relationships between rattle amplitude (a proxy for recording distance) and time of day (linear regression: t = - 1.33, df = 6.60, p = 0.187) or treatment (linear regression: t = −1.656, df = 24.9, p = 0.112). This indicates that any variation in rattle entropy throughout the day or among the treatments was not due to focal squirrels being closer to or further from the microphone.

### Statistical Analyses

For statistical analyses, we used R (version 3.5.1; R Developmental Core Team, 2018) with the package lme4 (version 1.17; Bates et al., 2015) to fit linear mixed-effects models and lmerTest version 3.0 (Kuznetsova et al.; 2017) to assess the significance of these models. For the capture-induced stress experiment, we included rattle collection method (post-trap or opportunistic) as a fixed effect. We included squirrel ID as random effects because we analyzed multiple rattles from the same squirrels across multiple years. Wilson et al. (2015) found no effects of age, sex, or Julian date on the acoustic structure of rattles recorded from this same population; we thus excluded these variables from analysis. We also found no year effects for any of the acoustic parameters measured – we ran a linear mixed effects model for each acoustic parameter with year included as a fixed effect, and found no significant relationships (*duration:* t = 0.20, df = 241.74, p = 0.84; *mean frequency*: t = −0.81, df = 264.18, p = 0.42; *entropy*: t = 262.2, df = −0.88, p = 0.379).

To examine the effects of administration of exogenous GCs on the acoustic structure of rattles, we fit three separate linear mixed-effects models – one for each of the three acoustic response variables (duration, mean frequency, entropy). Each model included an interaction between treatment group and time since treatment consumption (both linear and quadratic terms) as fixed effects, and squirrel ID (n = 44) as a random effect. In order to include the rattles of unmanipulated squirrels in this model, we found the average time at which the GC-treated and control (fed) squirrels consumed their treatment (1015 h) and set that as time of treatment consumption for all unmanipulated squirrels (i.e. unfed squirrels). For example, a rattle emitted at 1030 h would have a “time since treatment” value of 900 s, and a rattle emitted at 1000 h would have a time since treatment value of −900 s. Time since treatment consumption was standardized (mean (time of day) = 0, SD = 1).

If elevated plasma GCs alter rattle acoustic structure, we expected that the effects of the GC treatment on rattle acoustic structure would be strongest shortly after treatment consumption, the time frame in which circulating GCs should be highest using this treatment paradigm (Breuner et al., 1998; van Kesteren et al., submitted). Thus, we included a non-linear (quadratic) term for time since treatment consumption and its interaction with treatment because we expected that the effects of the treatment would exhibit a non-linear relationship, peaking shortly after treatment and then declined throughout the remainder of the day.

## Results

### Effects of capture-induced stress on rattle acoustic structure

Capture-induced stress caused pronounced differences in rattle acoustic structure: post-trap rattles were longer, higher in frequency, and noisier than rattles collected opportunistically. The average duration of post-trap rattles (4.77 ± 2.25 (SD) s) was significantly longer than that of opportunistic rattles (2.93 ± 1.28 s), a 63% increase (t = 3.78, df = 209.41, p < 0.001, Fig. 1A). The average mean frequency of post-trap rattles (7269.53 ± 1180.76 hz) was slightly but significantly higher than that of opportunistic rattles (6971.753 ±1007.37 hz) as well, a 4.3% increase (t =2.82, df = 218.01, p = 0.005, Fig. 1B). And finally, the average entropy of post-trap rattles (0.754 ± 0.035) was slightly but significantly higher than that of opportunistic rattles (0.712 ± 0.047), a 5.9% increase (t = 4.14, df =78.52, p < 0.001, Fig. 1C).

**Figure 1:**
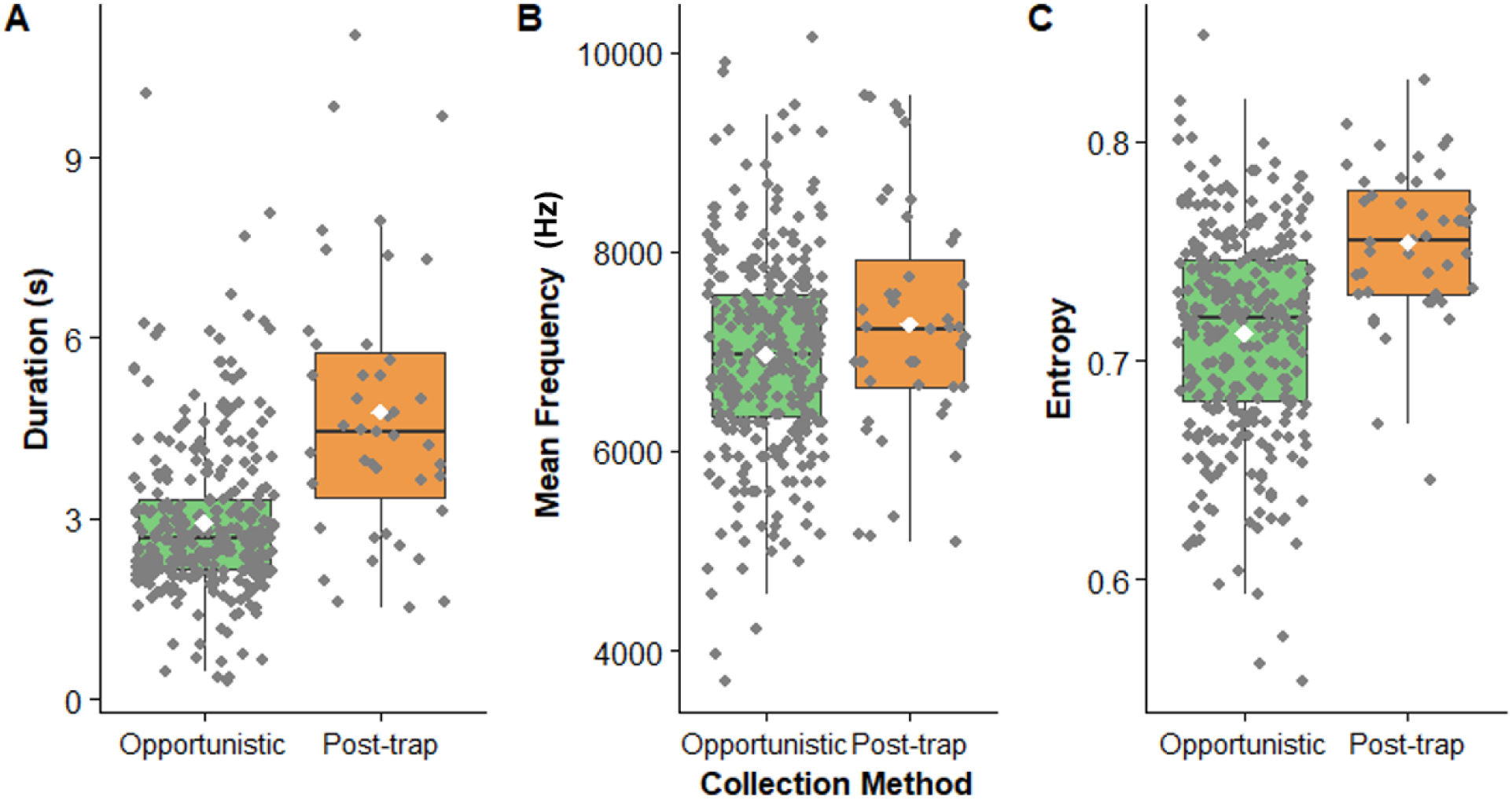
Effects of short-term stress (live-trapping and handling) on rattle A) duration (s), B) mean frequency (H**z),** and entropy. Post-trap rattles were collected within a minute of the squirrel exiting a trap and rattles collected opportunistically were collected from unprovoked squirrels. Post-trap rattles were significantly longer (t = 3.78, df **=** 209.41, p < 0.001, Fig. 1A), higher in frequency (t = 2.82, df = 218.01, p = 0.005, Fig. 1B), and higher in entropy (t = 4.14, df =78.52, p < 0.001, Fig. 1C). The black lines denote median, the white diamonds denote mean.

### Effects of administration of glucocorticoids on rattle acoustic structure

Administration of exogenous GCs did not produce the same effects on rattle acoustic structure as capture-induced stress; the rattle acoustic features of GC treated squirrels did not follow the predicted pattern of peaking after treatment and then declining as a function of time since treatment (Tables S1-S3, Fig. 2). There was, however, a significant linear interaction between treatment and the amount of time elapsed since treatment consumption on rattle duration (F_2, 677.4_ = 3.78, P = 0.02). This effect was largely driven by the increases in rattle duration observed in unmanipulated squirrels (Fig. 2A): rattles from unmanipulated squirrels increased in length throughout the day compared to those treated with GCs (b = 0.33, t = 2.67, P = 0.01, Table S1, Fig. 2A). Rattle durations of squirrels treated with GCs did not change differentially over the course of the day when compared with rattle durations of squirrels fed peanut butter only (b = 0.07, t = 0.73, P = 0.47, Table S1, Fig. 2A).

**Figure 2:**
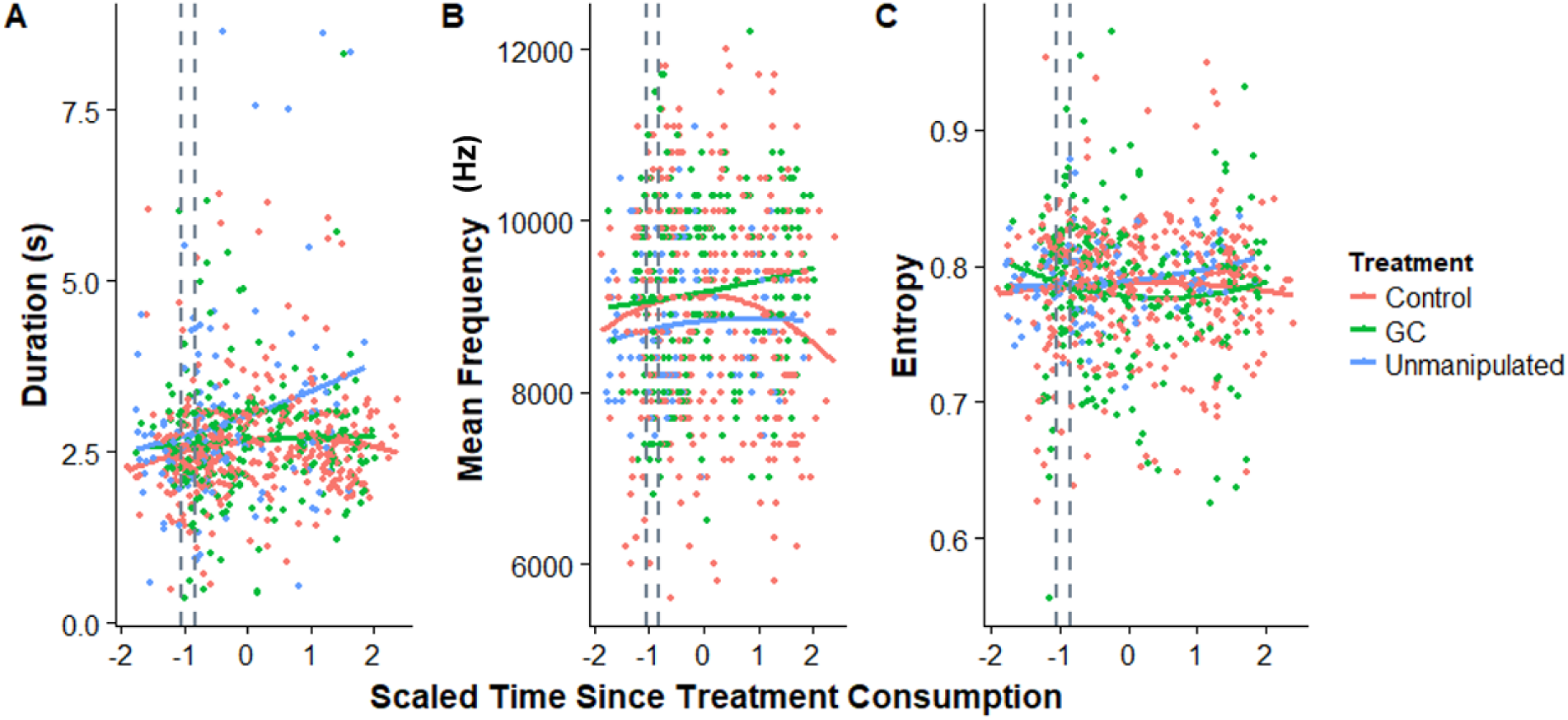
Effects of exogenous GCs (“GC”) and supplemental food (“Control”) on rattle A) duration (s), B) mean frequency (Hz), and C) entropy as a function of time since treatment. For unmanipulated squirrels, “time of treatment” is standardized at 1015 h, the average time of treatment consumption of GC-treated and control (fed) squirrels. The vertical gray dotted lines indicate the 1-hour time frame in which squirrels consumed their treatments. Squirrels fed supplemental food, exogenous GCs, or those that were unmanipulated had similar acoustic structure except that unmanipulated squirrels had significantly longer rattles than GC-treated squirrels as the time since treatment consumption increased (Table 1B). Time since treatment consumption was standardized (mean = 0, SD = 1).

There were no treatment effects on rattle mean frequency (F_2,2_ = 0.60, p = 0.63, Table S2) or entropy (F_2,56_ = 0.47, p = 0.63, Table S3) and the effects of the treatments on rattle mean frequency or entropy did not depend upon the amount of time that had elapsed since treatment consumption, as indicated by the lack of interactions between treatment and time elapsed since treatment consumption (both linear and quadratic terms). However, the mean frequency of rattles from squirrels recorded in all three treatment groups increased throughout the day (F_1,683.3_ = 4.77, p = 0.03). Overall, there were no significant non-linear effects of time since treatment consumption or its interaction with treatment on rattle duration, frequency, or entropy (Table S1, S2, S3).

## Discussion

Our study shows that short-term stress, in this case induced by live-capture and handling, significantly influences the acoustic structure of territorial vocalizations in red squirrels. Squirrels experiencing capture-induced stress produced rattles that were longer in duration, higher in frequency, and noisier (higher entropy) than rattles produced by control squirrels. However, we found little support for our second hypothesis that the changes in rattle acoustic structure induced by trapping and handling were caused primarily by increases in circulating GCs, despite the fact that GCs increase in response to trapping and handling. In our second experiment, the rattles of squirrels treated with GCs did not exhibit the expected structural distinctions from the rattles of control or unmanipulated squirrels over the treatment period.

The effects of short-term stress (trapping and handling) on rattle acoustic structure that we observed (longer duration, higher mean frequency, and higher entropy) are congruent with such trends in acoustic structure in relation to stress in many species. Chimpanzee screams, for example, increase in duration with the severity of an agonistic encounter (Slocombe et al., 2009). In dog barks (*Canis lupus familiaris,* Tokuda, 2002), human infant cries (Facchini et al., 2005), baboon grunts (*Papio hamadrayas,* Rendall, 2003), and meerkat alarm calls (*Suricata suricatta,* Manser, 2001), noisiness (entropy) increases with short-term stress. In many species, an increase in short-term stress is associated with an increase in pitch related characteristics. For example, during capture-release events, female bottlenose dolphins with dependent calves produce whistles of elevated frequency (*Tursiops truncatus,* Esch, 2009). The same pattern is observed in adult female African elephants (*Loxondota africana,* Soltis et al., 2005b), tree shrews (*Tupaia belangeri,* Schehka and Zimmerman, 2009), and Zebra finches (Perez et al., 2012): short-term stress is associated with an increase in vocalization pitch. In giant panda cubs (*Ailuropoda melanoleuca*), increased stress is associated with all of the trends in acoustic structure that we observed in post-trap rattles: longer duration, higher frequency, and increased noise (Stoeger et al., 2012).

Our results resemble those of Perez et al. (2012), who investigated how an environmental stressor (social isolation) and treatment with exogenous GCs affected vocalization structure in Zebra Finches. In their study, social isolation induced vocalizations of increased duration and pitch, and reduced overall vocal activity. However, oral administration of GCs only resulted in vocalizations with increased pitch, but no other effects were observed (Perez et al. 2012). The results from Perez et al. (2012) and our study suggest that short-term stressors alter vocalization structure but any increases in GCs caused by the short-term stressor are not solely responsible for the changes in vocalization structure.

Our findings and those of Perez et al. (2012) suggest that the acoustic structure of vocalizations can be altered by short-term stress, but the relationship between circulating GC levels and acoustic structure of vocalizations is not straightforward. Glucocorticoid treatment and capture-induced stress result in comparable concentrations of plasma GCs (capture: 400-1200 ng/ml, GC treatment: 500-1000 ng/ml; Dantzer, unpublished data), indicating that our GC treatment regime fairly accurately simulates the increase in plasma cortisol experienced as a result of capture. Thus, other hormones or neurochemicals may be implicated in modulation of the acoustic structure of vocalizations. For example, in rat pups, several classes of dopamine receptor agonists reduced the production of stress-induced ultrasonic vocalizations caused by isolation; this is a sign of reduced separation anxiety (Dastur et al., 1999). It is also possible that the acoustic structure of vocalizations has a non-monotonic dose response relationship with GCs. There is precedent for such a relationship: in white crowned sparrows, moderate doses of corticosterone induced elevated physical activity, whereas high levels did not (Breuner et al., 1998). We only provisioned squirrels with one dosage of GCs and so were unable to address whether lower or higher dosages of GCs would alter rattle acoustic structure. Together, this suggests the importance of considering additional mechanisms that may underlie the observed changes in vocalization acoustic structure.

Because treatment with exogenous GCs did not induce the same changes to vocalization structure as trapping, it is possible that these changes were produced by an effect of trapping besides stress. Because rattles function to advertise territorial ownership, it is possible that a squirrel that has been in a trap and unable to defend its territory for up to two hours, upon release, compensates by producing rattles that are longer and noisier. This hypothesis, however, would require explicit tests.

Our findings constitute further evidence that territorial vocalizations such as rattles contain more information than territorial ownership. In red squirrels, not only do rattles have the capacity to communicate stable information about the signaler’s individual identity and potential kin relationships (Digweed et al. 2012; Wilson et al. 2015; Shonfield et al. 2017), but also labile information, such as short-term stress. This layering of stable and labile encoded information in vocalizations is not uncommon, appearing across a number of animal taxa (Seyfarth and Cheney, 2003; Rendall, 2003; Blumstein and Munos, 2005; Soltis, 2005; Koren and Geffen, 2009; Terleph et al., 2016).

There are several hypotheses on the functional significance of these tendencies in vocalizations associated with high-stress contexts. In social species, the unpredictability hypothesis states that calls that contain more non-linearities are more difficult to habituate to, and thus noisy alarm calls are more likely to capture the attention of a conspecific in the event of a predatory or otherwise dangerous event (Blumstein and Recapet, 2009). Another hypothesis holds that screams produced when an animal is in imminent danger of predation serve to either startle and distract the predator, or solicit intervention from another animal, either a social group member, or a “pirate” predator that may attempt to steal the prey and unintentionally free it (Hogstedt, 1982). In the case of red squirrels, one hypothesis that can be envisaged is that honestly communicating stress to neighbors may advertise a willingness to aggressively defend one’s territory. Another possibility is that instead of honestly depicting a willingness to defend a territory, vocal cues of stress might inadvertently reveal that the caller faces some other challenge and might, therefore, be less capable of defending their territory. These two hypotheses, however, would need to be tested directly – for example, a playback study could test whether the rattles of stressed squirrels are more or less likely to deter territorial intrusions from neighboring squirrels than rattles of unstressed squirrels. If stress-influenced rattles are more likely to deter intruders, and if their production predicts an attack or further escalation by the signaler, then stressed rattles would be considered aggressive signals (Searcy and Beecher, 2009); if the opposite was the case, they would be considered index signals (Smith and Harper, 1995).

Though research on stress-induced changes to vocalizations has focused primarily on group-living species, the encoding of labile information such as short-term stress in vocalizations may have consequences in a population of solitary, territorial animals as well, perhaps enabling neighbours to eavesdrop on the physiological state of the signaler and adjust their own behavior or reproduction accordingly. Eavesdropping by conspecifics, or the acquisition of public information, may have important ecological consequences (Valone, 2007; Dall et al., 2010). For example, in many species, including red squirrels (Fisher et al., 2017; Lane et al., 2018), breeding earlier than other individuals in your population may be advantageous. Cues about the physiological state of a signaler contained in territorial vocalizations may provide an important source of information about when other individuals in the population are breeding – in red squirrels, the strongest level of selection for postnatal growth rate and birth date is the social neighborhood (Fisher et al., 2017). As such, labile information contained in vocalizations, such as stress state, may have broader ecological consequences by serving as public information and modifying the timing of reproduction in seasonally breeding species.

Overall, our results indicate that red squirrel territorial vocalizations contain labile information on physiological state, in addition to previously documented stable information about territorial ownership and individual identity. In some cases, this stable and labile information may interact – the stress state of the signaler might modify the ability of conspecifics to discriminate whether they are kin or non-kin, as proposed by Shonfield et al., (2017). These layers of encoded information may influence behavioral and reproductive dynamics, the effects of which merit exploration in future studies.

## Acknowledgements

We thank the Champagne and Aishihik First Nations for providing access to the land on which the study sites for this project were located, in particular Agnes MacDonald and her family for long-term access to her trapline. We also thank Zach Fogel and Noah Israel, the diligent field technicians whose work on the GC-induced experiment was crucial for its success. Funding was provided by the University of Michigan and National Science Foundation (IOS 1749627) to B. Dantzer as well as the Natural Sciences and Engineering Council of Canada (S. Boutin, M.M. Humphries, J.E. Lane, A.G. McAdam, D. Wilson). This is publication **XX** of the Kluane Red Squirrel Project.

## References

Andersson, M. 1994. Sexual Selection. Princeton, NJ: Princeton University Press.

August, P.V., & Anderson, J. G. T. 1987. Mammal sounds and motivation-structural rules: A test of the hypothesis. J. Mammal. 68:1–9.

Bates, D., Maechler, M., Bolker, B., & Walker, S. 2015. Fitting linear mixed-effects models using lme4. J.Stat. Soft. 67: 1-48.

Bayart, F., Hayashi, K.T., Faull, K.F., Barchas, J. D., & Levine, S. 1990. Influence of maternal proximity on behavioral and physiological responses to separation in infant rhesus monkeys (*Macaca mulatta*). Behav. Neurosci. 104: 98–107

Bee, M.A., Perrill, S.A., & Owen P.C. 1999. Size assessment in simulated territorial encounters between Male Green Frogs (*Rama clamitans*). Behav. Ecol. Sociobiol 45:177-184.

Beecher, M.D. 1989. Signaling systems for individual recognition: an information theory approach. Anim. Behav. 38: 248–261.

Beer, C. G. 1970. On the responses of laughing gull chicks (*Larus atricilla*) to the calls of adults II. Age changes and responses to different types of call. Anim. Behav. 18: 661–677.

Biben, M., Symes, D., & Masataka, N. 1986. Temporal and structural analysis of affiliative vocal exchanges in squirrel monkeys (*Saimiri sciureus*). Behaviour 98: 259-273.

Blumstein, D.T., & Munos, O. 2005. Individual, age and sex-specific information is contained in yellow-bellied marmot alarm calls. Anim. Behav. 69: 353–361.

Blumstein, D., Richardson, D., Cooley, L., Winternitz, J., & Daniel, J. 2008. The structure, meaning and function of yellow-bellied marmot pup screams. Anim. Behav. 76: 1055–1064.

Blumstein, D.T., & Récapet, C. 2009. The sound of arousal: the addition of novel non-linearities increases responsiveness in marmot alarm calls. Ethol. 115: 1074–1081.

Bosson, C.O., Islam, Z, & Boonstra, R. 2012. The impact of live trapping and trap model on the stress profiles of North American red squirrels. J. Zool. 288: 159-169.

Breuner, C., Greenberg, A., Wingfield, J. 1998. Noninvasive corticosterone treatment rapidly increases activity in Gambel’s White Crowned Sparrows (*Zonotrichia leucophrys gambelii*). Gen. Comp. Endocrinol. 111: 386-394.

Clark, A.P. 1993. Rank differences in the production of vocalizations by wild chimpanzees as a function of social context. Am. J. Primatol. 31: 159-197.

Dantzer, B., Newman, A.E.M., Boonstra, R., Palme, R., Boutin, S., Humphries, M.M., & Mcadam, A.G. 2013. Density triggers maternal hormones that increase adaptive offspring growth in a wild mammal. Science 340: 1215–1218.

Dastur, F.N., McGregor, I.S., & Brown, R.E. 1999. Dopaminergic modulation of rat pup ultrasonic vocalizations. Eur. J. Pharmacol. 382: 53–67.

Digweed, S., Fedigan, L., & Rendall, D. 2007. Who cares who calls? Selective responses to the lost calls of socially dominant group members in the white-faced capuchin (*Cebus Capucinus*). Am. J. Primatol. 69: 829-835.

Digweed, S., Rendall, D., & Imbeau, T. 2012. Who’s your neighbor? Acoustic cues to individual identity in red squirrel *(Tamiasciurus hudsonicus)* rattle calls. Curr. Zool. 58: 758–764.

Donald, J. L., Boutin, S., & Steele, M. A. 2011. Intraspecific cache pilferage by larder-hoarding red squirrels (*Tamiasciurus hudsonicus*). J. Mammal. 92: 1013–1020.

Ehret, G. 2005. Infant rodent ultrasounds - a gate to the understanding of sound communication. Behav. Genet. 35: 19–29.

Esch H.C., Sayigh, L.S., Blum, J.E., & Wells, R.S. 2009. Whistles as potential indicators of stress in bottlenose dolphins (*Tursiops truncatus*). J. Mammal. 90: 638–650.

Ey, E., Hammerschmidt, K., Seyfarth, R. M., & Fischer, J. 2007. Age and sex-related variations in clear calls of *Papio ursinus*. Int. J. Primatol. 28: 947-960.

Facchini, A., Bellieni, C.V., Marchettini, N., Pulselli, F.M. & Tiezi, E. B.P. 2005. Relating pain intensity of newborns to onset of nonlinear phenomena in cry recordings. Phys. Lett. A 338: 332–337.

Fisher, D., Boutin, S., Dantzer, B., Humphries, M., Lane, J., & McAdam, A. Multilevel and sex specific selection on competitive traits in North American red squirrels. Evolution. 71:1841-1854.

Fitch, W.T.S. 1997. Vocal tract length and formant frequency dispersion correlate with body size in rhesus macaques. J. Acoust. Soc. Am. 102: 1213–1222.

Fitch, W.T., Neubauer, J., & Herzel, H. 2002. Calls out of chaos: the adaptive significance of nonlinear phenomena in mammalian vocal production. Anim. Behav. 63: 407–418.

Fletcher, Q.E., Landry-Cuerrier, M., Boutin, S., Mcadam, A.G., Speakman, J.R., & Humphries, M.M., 2013. Reproductive timing and reliance on hoarded capital resources by lactating red squirrels. Oecologia 173: 1203–1215.

Smith, M.J. & Harper D.C.G. 1995. Animal Signals: Models and Terminology. J. Theor. Biol. 177: 305–311.

Hogstedt, G. 1982. Adaptation unto death: function of fear screams. Am. Nat. 121: 562–570.

Koren, L., & Geffen, E. 2009. Complex call in male rock hyrax (*Procavia capensis*): A multi-information distributing channel. Behav. Ecol. Sociobiol. 63: 581–590.

Koren, L., Mokady, O., & Geffen, E. 2008. Social status and cortisol levels in singing rock hyraxes. Horm. Behav. 54: 212–216.

Krebs, C.J., Boutin, S.A. & Boonstra, R. Ecosystem dynamics of the boreal forest: the Kluane project. New York, NY: Oxford University Press; 2001.

Kuztensova, A., Brockhoff, P., & Christianson, H.B. 2017. lmerTest Package: Tests in Linear Mixed Effects Models. J. Stat. Soft. 82: 1-26.

Lane, J.E., McAdam, A.G., McFarlane, S.E., Williams, C. T., Humphries, M.M., Coltman, D.W., Gorrell, J.C. and Boutin, S. 2018. Phenological shifts in North American red squirrels: disentangling the roles of phenotypic plasticity and microevolution. J. Evol. Biol. 31: 820-821.

Larsen, K.W. & Boutin, S. 1994. Movements, survival, and settlement of red squirrel (Tamiasciurus hudsonicus) offspring. Ecology. 75: 214-223.

Lingle, S., Rendall, D., & Pellis, S. M. 2007. Altruism and recognition in the antipredator defence of deer: 1. Species and individual variation in fawn distress calls. Anim. Behav., 73: 897–905.

Manser, M.B. 2001. The acoustic structure of suricates’ alarm calls varies with predator type and the level of response urgency. Proc. R. Soc. Lond. [Biol]. 268: 2315–24.

Marler, P. 1961. The Logical Analysis of Animal Communication. J. Theor. Biol. 1: 295–317.

Masataka, N., & Symmes, D. 1986. Effect of separation distance on isolation call structure in squirrel monkeys (*Saimiri sciureus*). Am. J. Primatol. 10: 271–278.

Mcadam, A., Boutin, S., Sykes, A., Humphries, M. Life histories of female red squirrels and their contributions to population growth and lifetime fitness. Ecoscience. 14: 362-369.

Morton, E.S. 1977. On the occurrence and significance of motivation-structural rules in some bird and mammal sounds. Am. Nat. 111: 855-869.

Muller, M.N., & Wrangham, R.W. 2004. Dominance, aggression and testosterone in wild chimpanzees: A test of the “challenge hypothesis.” Anim. Behav. 67: 113–123.

Nader, N., Chrousos, G.P., & Kino, T. 2010 Interactions of the circadian clock system and the HPA axis. Trends Endocrinol. Metab. 21: 277–286.

Perez, E.C., Elie, J. E., Soulage, C.O., Soula, H.A., Mathevon, N., & Vignal, C. 2012. The acoustic expression of stress in a songbird: does corticosterone drive isolation-induced modifications of zebra finch calls? Horm. Behav. 61: 573–581.

Perez, E.C., Mariette, M.M., Cochard, P., & Soulage, C. O. 2016. Corticosterone triggers high-pitched nestlings’ begging calls and affects parental behavior in the wild zebra finch. Behav. Ecol. 27: 1665–167.

Porges, W. 1995. Cardiac vagal tone: A physiological index of stress. Neurosci. Biobehav. Rev. 2: 225–233.

R Core Team. 2018. R: A language and environment for statistical computing. R Foundation for Statistical Computing, Vienna, Austria.

Reby, D., & McComb, K. 2003. Anatomical constraints generate honesty: acoustic cues to age and weight in the roars of red deer stags. Anim. Behav. 65: 519–530.

Rendall, D. 2003. Acoustic correlates of caller identity and affect intensity in the vowel-like grunt vocalizations of baboons. J. Acoust. Soc. Am. 113: 3390–3402

Searcy, W.A., & Beecher, M.D. 2009. Song as an aggressive signal in songbirds. *Animal* Behaviour, 78: 1281–1292.

Sacchi, R., Saino, N., & Galeotti, P. 2002. Features of begging calls reveal general condition and need of food of barn swallow *(Hirundo rustica)* nestlings. Behav. Ecol., 13: 268.

Sueur, J., Aubin, T., Simonis, C. 2008. Seewave, a free modular tool for sound analysis and synthesis. Bioacoustics.18: 213-226.

Schehka, S., & Zimmermann, E. 2009. Acoustic features to arousal and identity in disturbance calls of tree shrews (*Tupaia belangeri*). Behav Brain Res 203: 223–231.

Schmidt, K., Dall, S., & Van Gils, J. 2010. The ecology of information: an overview on the ecological significance of making informed decisions. Oikos 119: 304-316.

Seyfarth, R.M., & Cheney, D. L. 2003. Signalers and Receivers in Animal Communication. Annu. Rev. Psychol. 54: 145–173.

Shonfield, J., Gorrell, J.C., Coltman, D.W., & Boutin, S. 2017. Using playback of territorial calls to investigate mechanisms of kin discrimination in red squirrels. Behav. Ecol. 28: 1–9.

Siracusa, E., Morandini, M., Boutin, S., Humphries, M. M., Dantzer, B., Lane, J. E., & McAdam, A. G. 2017. Red squirrel territorial vocalizations deter intrusions by conspecific rivals. Behaviour 154: 1259-1273.

Slocombe, K.E., Townsend, S.W., & Zuberbühler, K. 2009. Wild chimpanzees (*Pan troglodytes schweinfurthii*) distinguish between different scream types: Evidence from a playback study. Animal Cogn. 12: 441–449.

Smith, C.C. 1968. The Adaptive Nature of Social Organization in the Genus of Three Squirrels *Tamiasciurus*. Ecol. Monograph. 38: 31-64.

Smith, C.C. 1978 Structure and function of the vocalizations of tree squirrels *(Tamiasciurus hudsonicus)*. J.Mammal. 59: 793-80.

Soltis, J., Leong, K., & Savage, A. 2005 African elephant vocal communication I: antiphonal calling behaviour among affiliated females. Anim. Behav. 70: 579-587.

Soltis, J., Leong, K., & Savage, A. 2005. African elephant vocal communication II: Rumble variation reflects the individual identity and emotional state of callers. Anim. Behav.70: 589–599.

Stoeger, A.S., Baotic, A., Li, D., & Charlton, B.D. 2012. Acoustic features indicate arousal in infant giant panda vocalisations. Ethol. 118: 896–905.

Studd, E.K., Boutin, S., McAdam, A.G., & Humphries, M.M. 2016. Nest attendance of lactating red squirrels (*Tamiasciurus hudsonicus*): Influences of biological and environmental correlates. J. Mammal. 97: 806–814.

Terleph, T.A., Malaivijitnond, S., & Reichard, U.H. 2016. Age related decline in female lar gibbon great call performance suggests that call features correlate with physical condition. BMC Evol. Biol. 16: 1–13.

Hothorn, T., Bretz, F., & Westfall, P 2008. Simultaneous Inference in General Parametric Models. Biom. J. 50: 346-363.

Valone, T. 2007. From Eavesdropping on Performance to Copying the Behavior of Others: A Review of Public Information Use. Behav. Ecol. Sociobiol. 62: 1-14.

Wilson, D.R., & Evans, C. S. 2012. Fowl communicate the size, speed and proximity of avian stimuli through graded structure in referential alarm calls. Anim. Behav. 83: 535–544.

Wilson, D.R., Goble, A.R., Boutin, S., Humphries, M.M., Coltman, D.W., Gorrell, J.C., Shonfield, J., McAdam, A. G. 2015. Red squirrels use territorial vocalizations for kin discrimination. Anim. Behav. 107: 79–85.

Van Kesteren, F., Delehanty, B., Westrick, S.W., Palme, R., Boonstra, R., Lane, J.E., Boutin, S., McAdam, A., & Dantzer, B. Experimental increases in glucocorticoids alter function of the neuroendocrine stress axis in wild red squirrels without negatively impacting survival and reproduction. Manuscript under review.

Yosida, S., & Okanoya, K. 2009. Naked mole-rat is sensitive to social hierarchy encoded in antiphonal vocalization. Ethol. 113: 113-131.

Zuberbuhler, K. 2009. Survivor Signals: The biology and psychology of animal alarm calling. Adv. Study Behav. 40: 277–322.

